# Reflections of Idiographic Long-Term Memory Characteristics In Resting-State Neuroimaging Data

**DOI:** 10.1101/2020.04.18.047662

**Authors:** Peiyun Zhou, Florian Sense, Hedderik van Rijn, Andrea Stocco

## Abstract

Translational applications of cognitive science depend on having predictive models at the individual, or *idiographic*, level. However, idiographic model parameters, such as working memory capacity, often need to be estimated from specific tasks, making them dependent on task-specific assumptions. Here, we explore the possibility that idiographic parameters reflect an individual’s biology and can be identified from task-free neuroimaging measures. To test this hypothesis, we correlated a reliable behavioral trait, the individual rate of forgetting in long-term memory, with a readily available task-free neuroimaging measure, the resting-state EEG spectrum. Using an established, adaptive fact-learning procedure, the rate of forgetting for verbal and visual materials was measured in a sample of 50 undergraduates from whom we also collected eyes-closed resting-state EEG data. Statistical analyses revealed that the individual rates of forgetting were significantly correlated across verbal and visual materials. Importantly, both rates correlated with resting-state power levels in the low (13-15 Hz) and upper (15-17 Hz) portion of the beta frequency bands. These correlations were particularly strong for visuospatial materials, were distributed over multiple fronto-parietal locations, and remained significant even after a correction for multiple comparisons (False Discovery Rate) and after robust correlation methods were applied. These results suggest that computational models could be individually tailored for prediction using idiographic parameter values derived from inexpensive, task-free imaging recordings.

## 1. Introduction

To provide a complete account of human behavior, psychological theories should explain behaviors at both the *nomothetic* level (that is, group aggregates and mean tendencies) and the *idiographic* level (that is, accurate characterizations of each participant; Allport, 1937), a claim that equally holds for computational cognitive models.

Although the nomothetic approach has been historically dominant and the majority of published computational models are fitted to group averages, the idiographic approach has been frequently advocated (Ritter & Gobet, 2000) and is often essential to translational applications of cognitive research. For instance, in intelligent tutoring systems, idiographic models are critical to providing appropriate adaptive feedback to specific errors and knowledge of individual students (Anderson et al., 1990).

In the idiographic approach, individuals can be characterized at the level of stable characteristics, or *traits*, or contingent situations, or *states*. An individual’s working memory capacity, for example, is relatively stable over time. On the other hand, a student’s domain knowledge is continuously expanded during studying and is thus better characterized as a *state*. Because they capture stable characteristics, traits are particularly useful to predict individual behaviors across tasks and over extended periods. An intuitive conceptualization is to think of idiographic traits as specific *values* of a model’s *parameter* (Collins, 2018; Daw, 2011; Lovett et al., 2000; Ritter & Gobet, 2000; Stocco, 2018). Working memory capacity, for example, can be captured by a parameter that represents the number of free slots in a buffer, and idiographic traits by different values of this parameter (Collins, 2018).

Idiographic trait parameters should exhibit at least two characteristics. First, they should have high stability over time. For instance, Sense et al. (2016) have shown that long-term memory decay is highly correlated across sessions administered on different days (*r* ∼ 0.8) and materials (*r* ∼ 0.5). Second, it should generalize *across tasks*: once a parameter has been estimated by fitting a model to an individual, the same parameter’s value should predict that individual’s performance in other tasks. For example, Maaß and colleagues (2019) were able to estimate the variability of the internal clock using a simple time production task (Maaß & van Rijn, 2018), and use it to predict performance in a more elaborated temporal reproduction task in a pre-clinical population.

In addition, we believe that a cognitive trait parameter should also be meaningfully related to brain features. An example of this is the correlation between procedural learning rate and the density of dopamine receptors (Stocco, 2018; Stocco et al., 2017). In this paper, we investigate whether individual variability in the long-term memory rate of forgetting is reflected in individual variability in electrophysiological measures of brain activity.

### 1.1 Resting-State Neuroimaging as a Window to Idiographic Parameters

The idea of correlating model parameters with brain activity is hardly new (see, for example, (Boehm et al., 2014; Schönberg et al., 2007; Tom et al., 2007). Most of these attempts, however, rely on *task-based* activity and thus suffer from a potential circularity: Because the model is designed to reproduce a specific type of task, the identified neural substrates are a function of the specific task assumptions as much as they are a function of the specific parameter.

This circularity can be circumvented by using *task-free* neuroimaging measures. These measures are made possible by the fact that spontaneous but organized brain activity exists even in the absence of any observable behavior (Fox et al., 2005). Task-free measures are recorded during “resting-state” sessions, in which participants refrain from doing anything: although they must remain awake, they are typically asked to just fixate a stimulus or close their eyes for a few minutes during which brain activity is recorded.

Although the nature and the functional significance of brain activity at rest are debated (Raichle, 2006; Raichle & Snyder, 2007), it is undeniable that resting-state brain activity carries signatures of measurable individual characteristics. For instance, resting-state fMRI can be used to classify individuals by age (Dosenbach et al., 2010), even among toddlers (Pruett et al., 2015), variations in resting-state correlations between regions have been linked to pathologies such as depression (Greicius et al., 2007) and Alzheimer’s disease (Sorg et al., 2007), and correlations between regions during resting-state can be used to predict behavioral performance during a task (Cole et al., 2016).

Although resting-state research typically focuses on fMRI, other modalities can be used, such as EEG. Instead of using correlations between time series (which are not meaningful for oscillatory signals), resting-state EEG data is first decomposed into different frequency components, and then the amount of variance of the whole signal that is explained by each frequency is estimated. The amount of variance is referred to as the frequency’s power and the power distribution over frequencies as the power spectrum. EEG power spectra are known be reliable and stable over time (McEvoy et al., 2000; Rogers et al., 2016) and contain sufficient inter-individual variability to be used to uniquely identify different participants, much like an electrophysiological fingerprint (Ma et al., 2015; Mohammadi et al., 2006; Näpflin et al., 2007). Finally, different features of the spectrum correlate with cognitive traits like intelligence (Doppelmayr et al., 2005) and language aptitude (Prat et al., 2016).

### 1.2. Idiographic Parameters in a Model of Long-Term Memory Forgetting Rate

Surprisingly, despite previous successes in identifying correlations between cognitive constructs (such as intelligence or aptitude) with resting-state EEG, no study to date has related task-free activity with the specific parameters of a cognitive model. The goal of this study is to demonstrate the possibility of measuring one specific and well-understood idiographic parameter from specific signatures of EEG data. Specifically, we will focus our attention on decay rate in long-term memory, a parameter that has an established tradition of use in cognitive models (J. R. Anderson, 1990; Shiffrin & Steyvers, 1997) and whose stability over time and cross-task reliability has already been previously studied (Sense et al., 2016).

The implementation used herein was inspired by Pavlik and Anderson (2008) and rooted in Anderson’s Bayesian account of memory (Anderson, 1990; Anderson & Schooler, 1991). A memory’s availability is proportional to a scalar meta-quantity, the *base-level activation*, which is the sum of the decaying traces of its previous usage. The rate of decay is determined for each memory trace separately and is a function of the *activation* of earlier encoded traces of the same memory when the memory is encoded and a memory specific intercept (see Van Rijn et al, 2009 and Sense et al, 2016 for details). The average of this memory specific decay rate intercept, *α,* is highly stable when the same individual is tested at different moments (*r* ∼ .80) and even across different materials (*r* ∼ .50), and thus meets the two first criteria for individual trait characteristics.

To measure α, Sense et al. (2016) asked participants to memorize a series of paired associates relating an old, familiar item with an entirely new item using a previously developed, adaptive memory testing procedure (Van Rijn et al., 2009). In one of their studies, the cues were (unfamiliar) Swahili words, with their (familiar) Dutch translation as the answers.

### 1.3 Research Question and Considerations About Effect Sizes

In this study, we replicated the experimental setup of Sense et al. (2016) and asked participants to perform an iterative paired-associates test, using both verbal (i.e., Swahili words) and non-verbal materials (i.e., maps of locations in the United States). The estimated mean value of α for each individual was then correlated with features of an individual power spectrum, estimated from a 5-minute resting-state EEG recording collected before the behavioral task. We expected to find a correlation between the value of the decay rate α and the spectral power in frequency bands, such as the alpha and beta rhythms, that have been previously correlated with similar memory-related constructs such as language learning rate (Prat et al., 2016).

## 2. Materials & Methods

### 2.1 Sample Size

This study’s goal is to identify a correlation between two different measures, one at the behavioral level (forgetting rate) and one at the neurophysiological level (spectral power), that purportedly target the same underlying biological process (i.e., the reduction in the availability of memory traces). The expected size of the correlation is affected by the stability of both measures. Sense et al (2016) estimated that the stability for α across sessions (measured as test-retest correlation) is *r* ∼.80. Unpublished data from our laboratory show that the test-retest correlation of EEG power spectra, using the very same equipment reported in this study, is approximately *r* ∼.60 across all channels and frequency bands (Prat et al., in preparation). Thus, a conservative upper bound for the maximum reliable correlation that we can expect is .80 × .60 = .48. Given this effect size, we estimated that collecting data from *N* = 50 participants would give us sufficient power to avoid a Type II error with probability *β* = 0.95 with an uncorrected significance level of *p* < 0.05.

### 2.2 Participants

Fifty-three native English speakers (32 females) aged between 18 to 33 years old (mean = 21) were recruited from the undergraduate population at the University of Washington, Seattle. None of the participants reported any familiarity with the Swahili language. Only native English speakers were selected to ensure that their performance on the Swahili vocabulary test (which required memorizing pairs of English-Swahili words) was not confounded by significant differences in English proficiency and English vocabulary exposure. Data from three participants were excluded because of either equipment failure (one male participant) or too few data points (< 75 artifact-free epochs in each channel: two females), leaving 50 subjects’ data (30 females) in the final analysis.

### 2.3 Materials

#### 2.3.1 Language Background

Participants’ language background was measured using the Language Experience and Proficiency Questionnaire (LEAP-Q: Marian, Blumenfeld, & Kaushanskaya, 2007). The LEAP-Q reports were used to ensure that all participants were native English speakers (Average age of English acquisition: 14 months old; 100% in self-rated English speaking proficiency; 100% in self-rated English understanding proficiency) and none of them had been previously exposed to Swahili.

#### 2.3.2 Swahili Vocabulary Learning Task

Twenty-five Swahili-English word pairs were selected from a previous study (Van den Broek, Segers, Takashima, and Verhoeven, 2014). The task dynamically interleaved study trials and test trials. On study trials, a Swahili word (e.g., “Samaki”) and its corresponding English equivalent (“fish”) were presented simultaneously on the screen. On test trials, only a Swahili word was presented on the screen as cue (e.g., “Samaki”), and participants were asked to respond by typing the corresponding English word (“fish”) in an empty textbox. Trials are self-paced, with no cap on the time allowed for a response. Consecutive trials are separated by a 600ms ISI after a correct response and a 4,000ms ISI after feedback following an incorrect response. The order of repetitions and moment of introduction for each item were determined by the adaptive scheduling algorithm outlined below and described in more detail in Van Rijn et al. (2009), van der Velde et al. (2020), and Sense et al. (2015; 2016). The adaptive algorithm schedules the presentation of future test trials based on internal estimates of the corresponding pair’s memory activation and decay rate (see Section 2.4.2 for details), and presents new study trials if no test trial is scheduled. Participants received corrective feedback after each response. A running timer and the ongoing response accuracy remained visible on the screen during each session. The task lasted approximately 12 minutes.

#### 2.3.3 Map Learning Task

The map learning task used the same procedure and scheduling algorithm as the vocabulary learning task but different stimuli. Instead of a Swahili word with the corresponding English translation, the stimuli consisted of an outline of a US map with dots representing twenty-five real but small cities (e.g., Buchanan, IL), each with a unique name. On each study trial, participants were shown a map with one location highlighted in red and were instructed to learn the corresponding city name, which was displayed along with the map. On test trials, participants were shown a map with one highlighted location and were asked to type in the city name. All other details were identical to the Swahili vocabulary task.

### 2.4 Procedures

Participants were asked to complete the LEAP-Q and a demographic survey before a five-minute eyes-closed and five-minute eyes-open resting EEG recording were collected. Then, they completed both learning tasks with the order of learning tasks counterbalanced based on the parity of participant IDs. At the end of the session, participants filled out a survey about perceived difficulty per task, data of which are not analyzed in this manuscript.

#### 2.4.1 Acquisition and Preprocessing of EEG data

Two continuous 5-minute recordings of resting-state EEG data were collected for each participant using a wireless 16-channel headset (first-generation Emotiv EPOC, Australia) with a sampling rate of 128 Hz. The reference channels were the DMS and CRL electrodes over the parietal lobe. During the *eyes-closed* resting-state EEG recording, participants were instructed to close their eyes, clear their mind, and relax, all while in a dark room. During the *eyes-open* resting-state EEG recording session, the experimenter turned on the light in the room and instructed participants to relax and look at a black fixation point on a white screen for 5 minutes while their EEG data were recorded. Although eyes-closed recordings seem to be slightly more common in recent literature (e.g., Benz et al., 2014; Prat et al., 2016, 2019), good arguments can be made in favor of eyes-open recordings. For example, eyes-open recordings might provide a better baseline to understand mental processes that include visual processing (Barry et al., 2007) and their smaller alpha peak is less likely to contaminate the theta and beta frequency bands (Klimesch, 1999). For these reasons, we decided to collect and compare both types of recordings.

The eyes-opened and eyes-closed EEG data were processed separately using the following procedure. Each channel’s time series was divided into two-second epochs with a 0.5-second overlap. Epochs containing significant artifacts (e.g., eye blinks, excessive motion, or signal deflections greater than 200 μV) were excluded from the analysis. All remaining 2-second epochs underwent Fast Fourier Transformation (FFT). The spectra of each epoch were then averaged together for each channel.

Note that, because the FFT algorithm was applied epoch-wise, the resolution of the spectrum was constrained by having an upper value of 64 Hz (half of the sampling rate, per Nyqvist’s theorem) and frequency resolution of 0.5 Hz (corresponding to the reciprocal of the epoch’s duration, that is, 2 s). Because the average spectrum for the entire time series was obtained by averaging the spectra of the individual epochs, it also had a resolution of 0.5 Hz and extended from 0 to 64 Hz.

Finally, the spectral powers within each frequency band were averaged to yield mean power values in the theta (4-8Hz), alpha (8-13Hz), beta (13-30Hz) bands. The beta was further subdivided into low beta (13-15Hz), upper beta (15-18Hz), and high beta (18-30Hz) bands. This was done, in agreement with previous studies (Prat et al., 2016, 2019; Zhou et al., 2020), to allow for a better resolution of the power spectra in the beta band, which spans a larger set of frequencies than theta and alpha and has been previously found to be related to long-term memory encoding and retrieval (Hanslmayr et al., 2012; Sederberg et al., 2007). The signal-to-noise ratio afforded by our equipment did not permit to reliably detect signals at higher frequencies, while very low frequencies tend to be contaminated by sporadic and non-physiological sources of noise (Cohen, 2014). Due to these considerations, gamma (> 30 Hz) and delta (0-4 Hz) frequency bands were excluded from the analysis. Any channel with fewer than 75 artifacts-free epochs for each individual was excluded, resulting in the removal of one channel (0.14% of the data) in the final analysis. This preprocessing pipeline was the same used in a previous study on neural markers of language learning (Prat et al., 2016; 2019). The analysis script can be found at http://github.com/UWCCDL/QEEG.

#### 2.4.2 Analysis of Behavioral Data

The main behavioral measure that will be correlated with EEG power is the average rate of forgetting α, which was estimated separately for each learning session (maps vs. vocabulary) of each participant. Here, we employed the same approach as in Sense, Behrens, Meijer, and van Rijn (2016): The parameter is estimated for each item separately, and an individual session’s parameter is computed by averaging the final values of all items that have been repeated at least three times. The aggregate value thus reflects the mean individual rate of forgetting and it reflects how quickly, on average, each participant forgot each type of material.

##### Memory Decay Model

The estimation of α was based on the formal model of memory decay (Anderson, 1990; Pavlik & Anderson, 2005; Sense et al., 2019) and the model of paired associates learning established by (Anderson, 1974). It assumes that, during the task, participants learn by forming a new memory *m* that links an existing memory (e.g., the word “fish”) to a new element (e.g., the word “samaki” in Swahili). In response to a study probe, participants use the newly learned word (“samaki”) as a cue to retrieve *m* (“samaki” - “fish”); if the retrieval succeeds, participants can type in the correct answer (“fish”). In this model, the key factor determining performance is the availability of memory *m*, which is related to its activation *A*(*m*). Every time *m* is retrieved or re-encoded, a new trace *i* is created. The final activation of *m* is the sum of the decaying activation of all its traces:

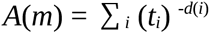

where *t_i_* is the time elapsed since the creation of *i* and *d*(*i*) is the decay rate of trace *i*. The decay rate is related to the *m*’s activation at the moment each trace is created, thus capturing the fact that traces with higher initial activation decay faster than traces with lower activation (Pavlik & Anderson, 2005). Specifically:

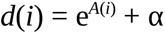

with *A*(*i*) being the activation of m at the moment *i* was encoded, and *c* being a constant scaling value. Thus, if the times at which a memory *m* has been encoded, re-encoded, and retrieved are known and controlled for, its activation depends only on the rate of forgetting α. In turn, α can be estimated indirectly by examining participants’ response latencies (i.e., times to typing the first key in response to study probes), which are related to activation by the equation:

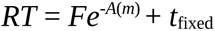

with *F* being a group-level free parameter and *t*_fixed_ representing a fixed offset for non-memory related processes.

##### Adaptive Stimulus Presentation

For the current study, a variant of the described memory decay model was used to adaptively devise repetition schedules while participants were studying the materials. Using the equations outlined above, the adaptive model fixes all but the α parameter. Each newly introduced word pair is initialized with a default α value of 0.3. Any time a new trial commences, the activation of all previously studied word pairs can be projected forwards in time. If any word pair has a predicted activation that is below a fixed threshold of - 0.8 at time *t* = 15 seconds from now, the word pair with the lowest activation is scheduled for repetition on the current trial; if not, a new word pair is randomly chosen and added to the rotation. This ensures that word pairs are scheduled for rehearsal before they are expected to be irretrievable and new items are only introduced once previous items are estimated to decay slowly. The α value for each word pair is updated after each collected response once it has been encountered three times: If the recorded RT differs more than 500*ms* from the predicted RT, a binary search is performed to find the α value that provides the best fit to the word pair’s recent history. More detailed descriptions of the adaptive system and its cognitive psychological motivations can be found in (Sense et al., 2016; van der Velde et al., 2020).

Since the scheduling of trials is adaptively driven by participants’ responses during study, the exact number of trials, repetitions per word pairs, order of their introduction, etc. will vary between participants. What is constant is the duration of the study session as well as the set of materials that are studied.

## 3. Results

### 3.1 Behavioral Results

Figure 1A shows the distribution of the rates of forgetting for the two types of materials, estimated for each participant, and plotted against each other. For the Swahili items, the mean rate of forgetting is α = 0.283 (*SD* = 0.062) and for the US Maps, the mean rate of forgetting is α = 0.336 (*SD* = 0.051). The figure shows that a participant’s rate of forgetting for one type of material is highly correlated with the rate of forgetting for the other type of material [*r*(50) = 0.59; *t*(48) = 5.08; *p* < 0.001). This correlation is comparable to the one reported in the study these materials were adapted from (approx. 0.50–0.55, see Table 1 in Sense et al., 2016). The figure also shows that the rates of forgetting tend to fall above the diagonal equality line, indicating that the rates of forgetting were generally higher for the US maps (range: 0.21–0.47) than for the Swahili words (range: 0.12–0.39; paired *t*(48) = 7.24, *p* < 0.001).

**Figure 1:**
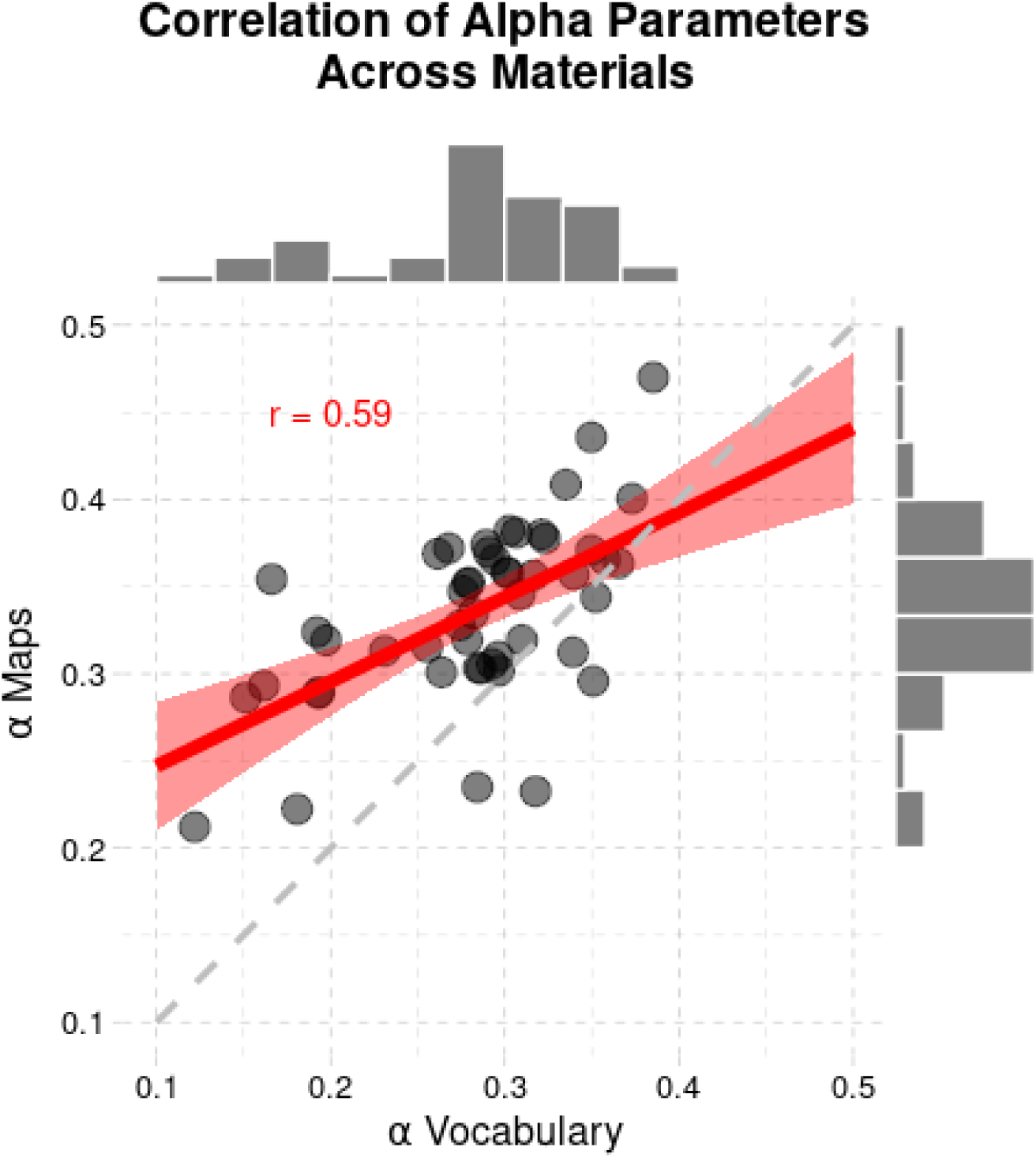
Average rates of forgetting for verbal (vocabulary) and visuospatial (US maps) materials plotted against each other. The dashed grey line represents the identity line; the thick red line represents the regression line, and the red shaded area represents the 95% confidence interval around the regression estimate.

The distribution of the estimated rates of forgetting in Figure 1 shows the same general pattern that was observed in other contexts, both in terms of the spread of the values as well as their relative distributions. As in earlier studies, the vocabulary materials are estimated to be easier than the maps (i.e., lower rates of forgetting), and rates of forgetting usually have values between roughly 0.1 and 0.4. For example, Figures 4 and 5 in Sense, Behrens, Meijer, and van Rijn (2016) and the scatterplot in Figure 2 of Sense et al. (2018) suggests that the distribution of the rates of forgetting in our sample is similar to what was found in previous studies, with values centered around 0.3, tapering off towards 0.2 and 0.4, with some individuals reaching rates of forgetting lower than 0.2.

**Figure 2:**
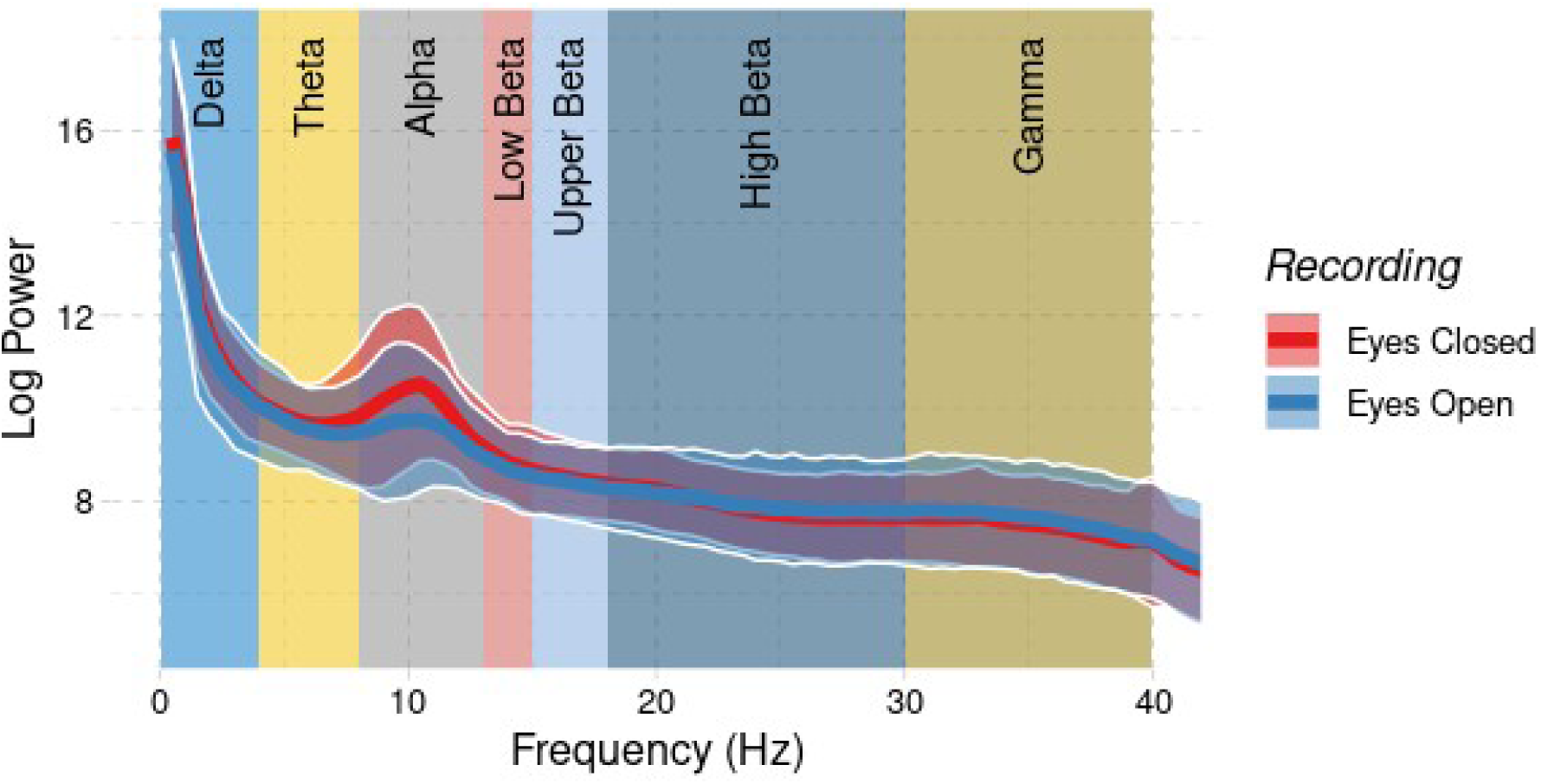
Power spectra averaged across all participants and channels, with the different recordings indicated in red (eyes closed) and blue (eyes open). The thick colored line at the center of the ribbon represents the mean power at every frequency, the shaded ribbon represents the standard deviation for the corresponding recording, and the relevant frequency bands are represented as different background colors.

**Figure 3:**
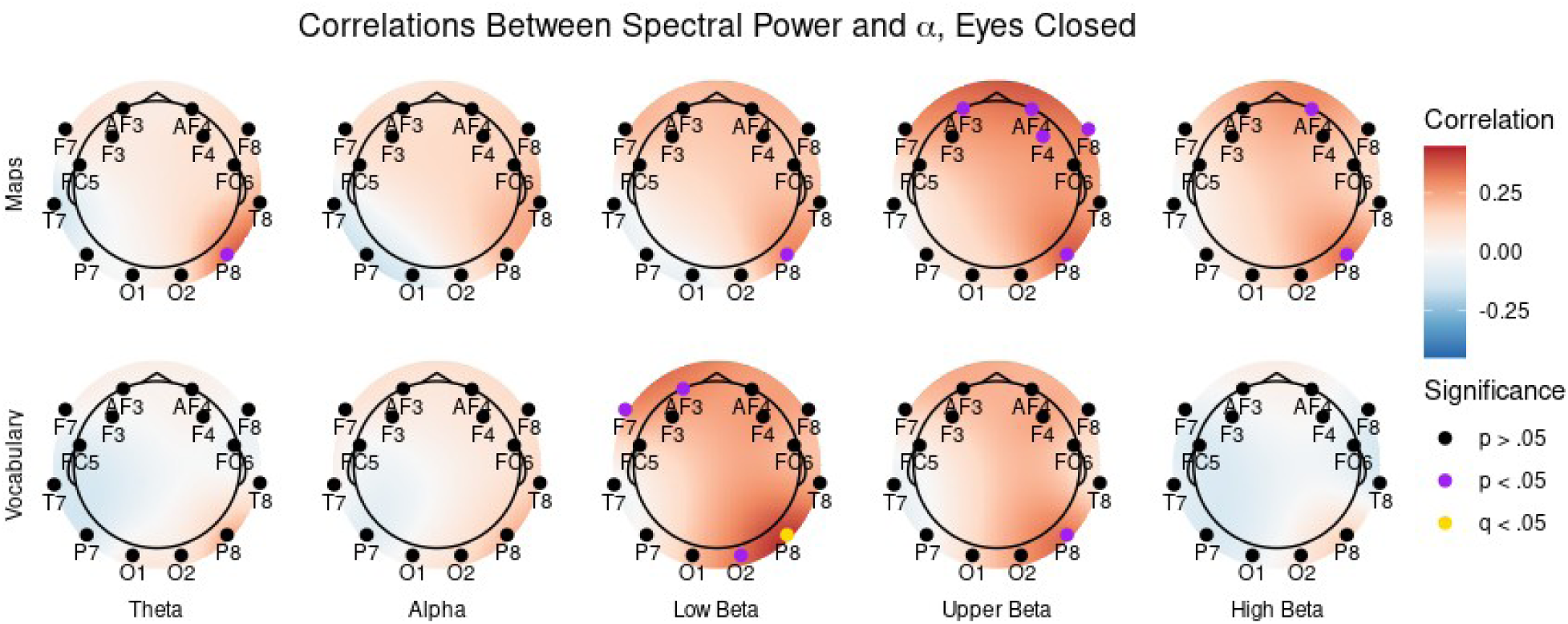
Topological maps of the correlations between mean power in each relevant frequency band (Theta, Alpha, and Low, Upper, and High Beta) and the individual values of forgetting rate for verbal (Swahili vocabulary, top) and visuospatial materials (US Maps, bottom) during eyes-closed EEG recordings. * p < .05; ** q < .05

**Figure 4:**
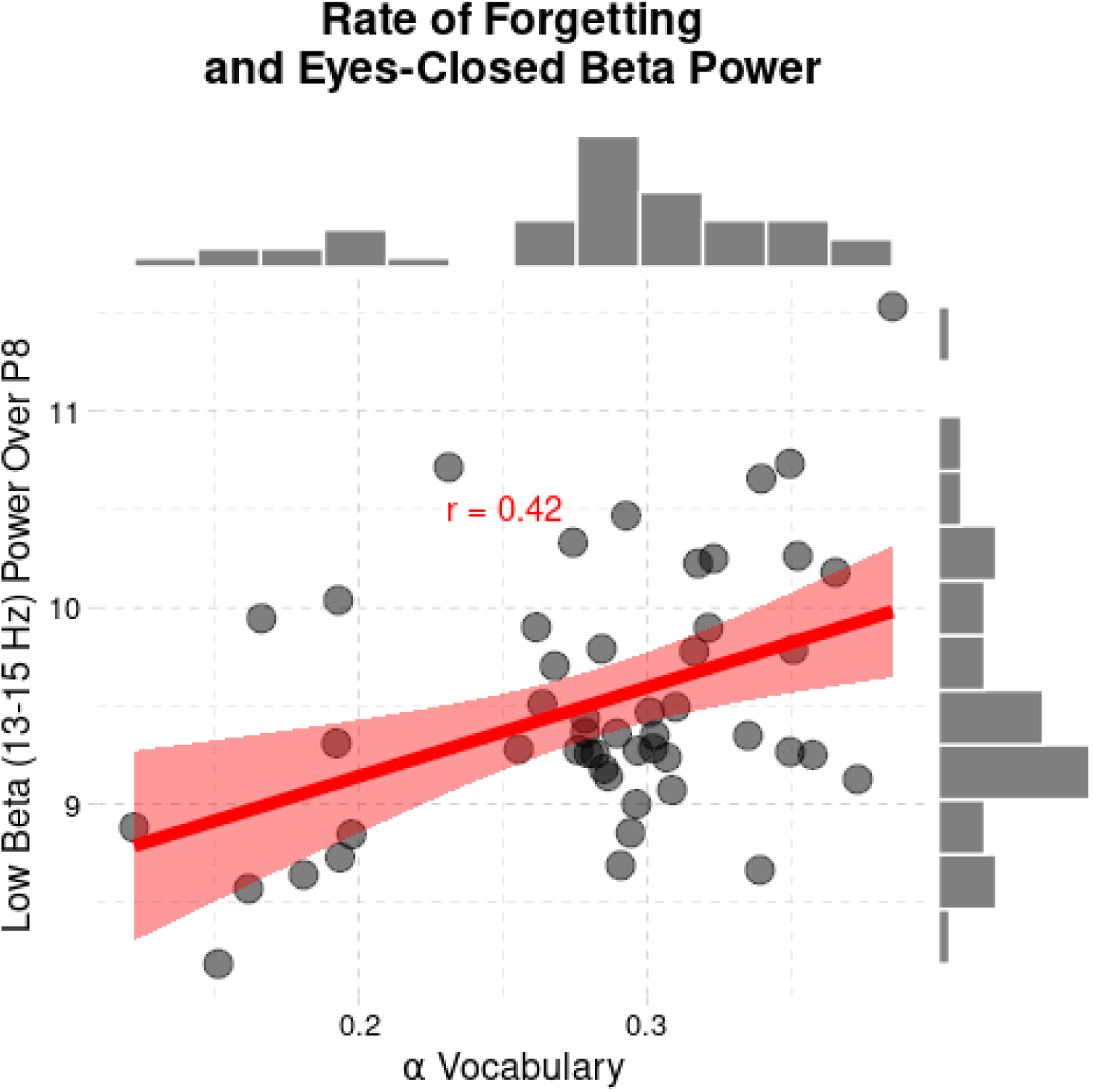
Correlation between rates of forgetting for the Swahili vocabulary task and mean low-beta power over P8.

**Figure 5:**
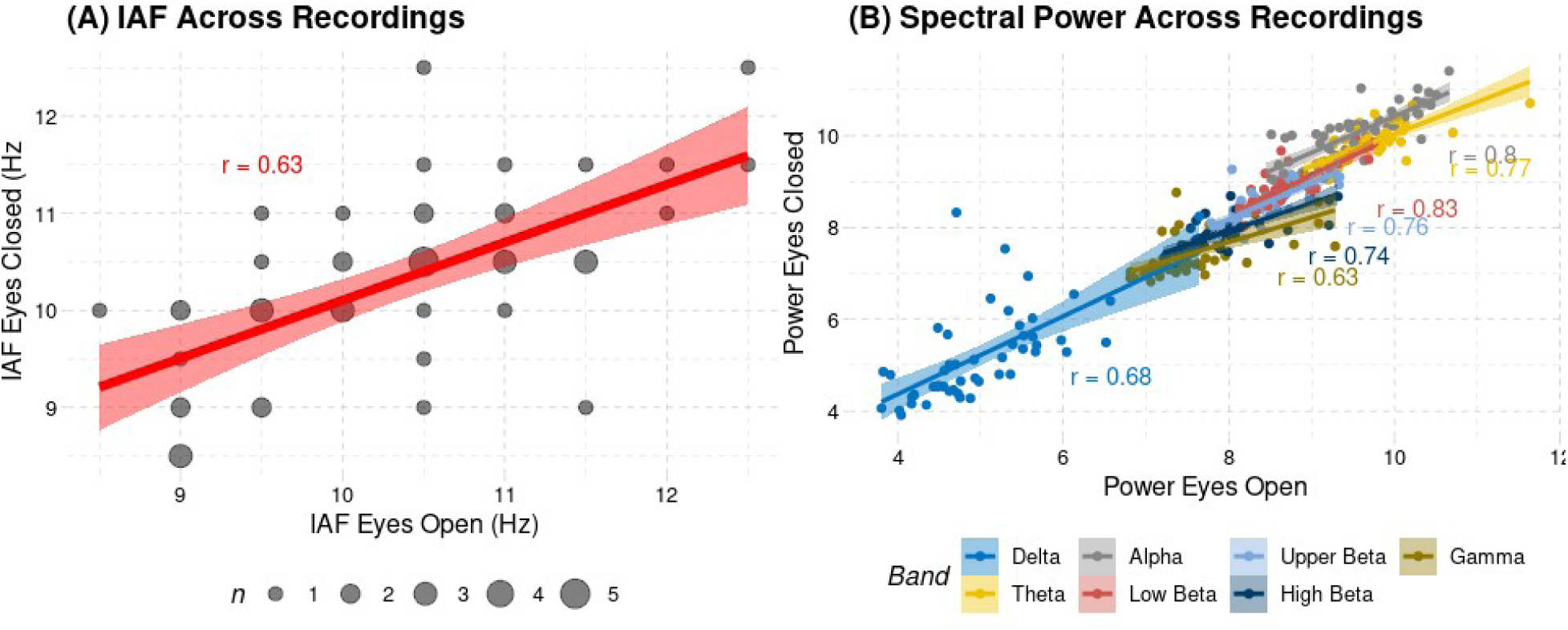
(A) Correlation between the individual alpha frequency (IAF) observed in eyes-closed vs. eyes-open EEG recordings. (B) Correlation between spectral power in different frequency bands in eyes-closed vs. eyes-open recordings.

### 3.2 EEG Results

The EEG power spectrum was extracted for each channel and visually inspected to ensure its consistency across channels and its conformity to the established distribution of power across frequency bands. Figure 2 depicts the power spectrum over frequencies in the 0-40 Hz range averaged across all 14 channels, with the seven relevant frequency bands marked by different background colors. Although the gamma and delta bands were not included in the analysis, they are depicted in Figure 2 for completeness. As expected, all channels exhibit the same power spectral profile, with the characteristic 1/*f* power distribution that is typical of electrophysiological signals (Buzsaki, 2006; Cohen, 2014). The peak in the alpha band (8-13 Hz), known as the individual alpha frequency (IAF), identifies the center of the alpha frequency band; although some authors prefer to use a priori defined ranges for each frequency (Prat et al., 2016), others prefer to define the alpha band on the bases of the mean IAF across participants and offset the other bands in relation to it (Doppelmayr et al., 2002). In our data, the mean IAF was 10.5 Hz in all channels (Figure 2). This value falls exactly at the center of the predefined range, thus making our results compatible with both approaches and implying that the predefined frequency bands are also representative of our sample.

### 3.2 Eyes-Closed EEG Power and Forgetting Rate

Next, we examined the correlations between the rates of forgetting and the power across channels in each of the following frequency bands: theta (4-8 Hz), alpha (8-13 Hz), beta (13-30 Hz), low beta (13-15 Hz), mid-beta (15-18 Hz), and high beta (18-30 Hz). In the following sections, we report all the correlations significant at *p* < .05, corresponding to a Pearson correlation value of *r* > 0.27. Because of the number of correlations that were computed, we also applied the False Discovery Rate procedure (Benjamini & Hochberg, 1995) to account for multiple comparisons, and highlight results that pass the corrected threshold of *q* < .05.

#### 3.2.1 Rates of Forgetting for Verbal Materials

The top row of Figure 3 presents the correlations between power at specific channel locations and the rate of forgetting in the Swahili word learning task across frequency bands. Significant positive correlations were found in the low and upper beta frequencies (13–18Hz) over the right parietal lobe [P8 in the 10–20 system, *r*(50) > .35, *p* < .02]. The largest positive correlation [*r*(50) = .42, *p* = .002] was observed at the right parietal channel (P8 in the 10–20 system) in the low beta band (13–15 Hz). This particular location and power band were also the only ones to show a significant effect when correcting for multiple comparisons (*q* = .03; Figure 4). Significant positive correlations that did not survive corrections for multiple comparisons were found for the low beta band at the bilateral frontal regions [AF3, F4, F7, *r*(50) > .27, *p* < 0.05, *q* > .15] and right occipital region [O2, *r*(50) = .34, *p* = .02, *q* = .11], and for mid-beta power at the right parietal lobe [P8, *r*(50) = .34, *p* = .02, *q* = .24].

#### 3.2.2 Rates of Forgetting for Visuospatial Materials

As illustrated in Figure 3 (bottom row), the correlations between EEG power and rates of forgetting in the map learning task showed a similar pattern of results; in fact, the two sets of correlations were themselves correlated at *r*(70) = .61 (*p* < .0001).

However, the correlations between rate and forgetting and spectral power were overall smaller in the case of visuospatial materials. As in the case of the vocabulary task, no negative correlation reached significance at *p* < .05. No positive correlation was significant at the corrected value of *q* < .05; however, significant positive uncorrected correlations between the rate of forgetting and power in the beta frequency band were observed. Specifically, power in the low beta band was positively correlated with the rate of forgetting at the right parietal lobe [P8, *r*(50) = .30, *p* = .04, *q* = .32]. The significant positive correlation between power in the beta band and rate of forgetting was observed distributing at bilateral frontal [AF3, AF4, F4, *r*(30) = .29–.34, *p* < .04, *q* < .32], right frontal [F8, *r*(50) = .32, *p* = .02, *q* = .09], and right parietal regions [P8, *r*(50) = .36, *p* = .009, *q* = .09]. Power in the high beta (18-30Hz) was positively correlated with the rate of forgetting at the right frontal [AF4, *r*(50) = .28, *p* = .05, *q* = .68] and right parietal region. [P8, *r*(50) = 0.33, *p* = .02, *q* = .25].

### 3.3 Eyes-Open EEG Power and Forgetting Rate

We also examined the eyes-open recordings. As noted above, eyes-open recordings are slightly less common in the EEG literature, likely because the alpha peak is less prominent. However, they are also more directly comparable to fMRI recordings, which are typically acquired while participants keep their eyes open.

#### 3.3.1 Relationship To Eyes Closed Recordings

Because eyes-open recordings are less common, we first investigated the degree to which their large-scale characteristics are comparable to the more common eyes-closed recordings. To do so, we calculated the mean spectrogram for all channels of each participant and found them to be essentially identical to their eyes-closed counterparts, except for a diminished, but still recognizable, alpha peak. We also calculated the mean spectrogram across all channels and directly compared the mean eyes-closed and eyes-open spectrograms, as shown in Figure 2. As is apparent, the spectrograms are essentially overlapping, with the notable difference of the alpha peak being much more prominent, as expected, in the eyes-closed recordings. Furthermore, the portion of the frequency spectrum in which the eyes-closed recordings exhibit a higher power than the eyes-open recordings falls in the alpha band, confirming that the group boundaries of our frequency bands were correct. Because the alpha peak was still identifiable, albeit diminished, in individual channel recordings, we calculated the IAF for eyes-open recordings as well. The IAF was calculated for each channel, and a participant-level estimate was calculated by taking the modal value across channels. Our results showed that, once more, the mean IAF across participants was 10.4, once more in the middle of the 8-13 alpha range. The correlation between each participant’s individual IAF identified from eyes-closed and eyes-open recordings was *r*(50) = 0.63, *p* < .0001, with a mean standard deviation of just 0.82 Hz in the difference between the values of the IAF in the two recordings (Figure 5A). Finally, as the last control check, we also calculated the mean correlation between the observed spectral power for each frequency band across eyes closed and eyes open recordings; all the correlations were found to be significant (r > .63, p < .0001), especially in the cognitive bands of interest (theta, alpha, and the beta bands), in which the observed correlation values were between *r*(50) = .74 and *r*(50) = .83 (Figure 5B).

Taken together, the recordings suggest that our eyes-open largely reflected the same neural characteristics as the eyes-closed ones at the *group* level. However, It is possible that variance in the eyes-open recordings contain additional, sensitive information at the *individual* level, which might be lost when the data is averaged and could be critical for the type of investigation pursued herein. Hence, we proceeded to conduct the same analysis and found significant positive correlations between beta power and rates of forgetting for verbal and visual materials mostly concentrated in the bilateral frontal and right parietal sites, as shown in the topological maps of Figure 6.

**Figure 6:**
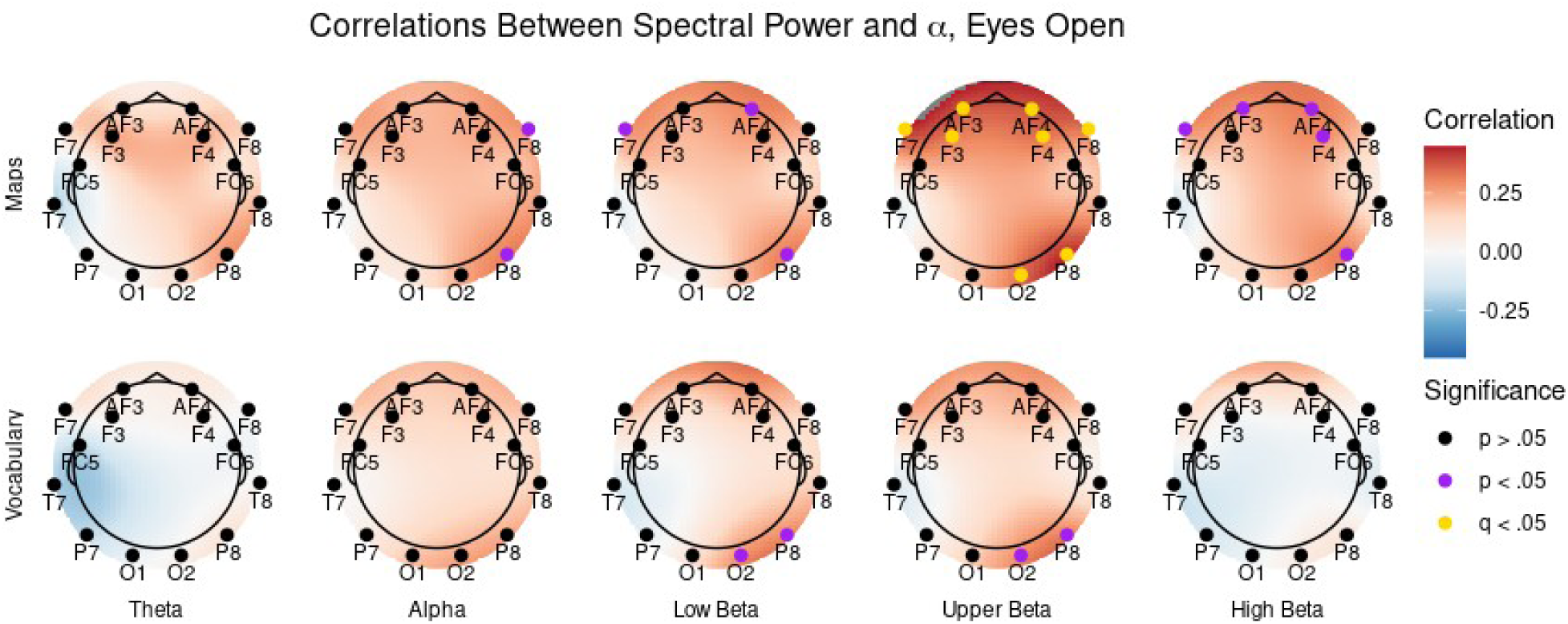
Topological maps of the correlations between mean power in each relevant frequency band (Theta, Alpha, and Low, Upper, and High Beta) and the individual values of forgetting rate for verbal (Swahili vocabulary, top) and visuospatial materials (US Maps, bottom) during eyes-open EEG recordings. * p < .05; ** q < .05

#### 3.3.2 Rates of Forgetting For Verbal Materials

In the case of verbal materials, the results were similar but somewhat weaker than in the case of eyes-closed recordings. None of the correlations survived the multiple comparisons corrections. Significant uncorrected positive correlations were found in the low beta band for P8 [*r*(50) = .28, *p* = .05, *q* = .28] and O2 [*r*(50) = .29, *p* = .04, *q* = .26] and in the upper beta band, again, for P8 [*r*(50) = .30, *p* = .04, *q* = .28] and O2 [*r*(50) = .31, *p* = .03, *q* = .27]. No significant negative correlation was found.

#### 3.3.3 Rates of Forgetting for Visuospatial Materials

As was the case in eyes closed recordings, the correlations between rate of forgetting for visuospatial materials and spectral power closely resembled those of verbal materials [*r*(70) = .81, *p* < .0001] Only, this time the correlations were stronger, with eight different channels showing significant positive correlations at the corrected threshold of *q* < .05. All of these correlations were found in the upper beta frequency band (13-15 Hz), and are illustrated in Figure 7. Specifically, they were found in the bilateral frontal channels AF3 [*r*(50) = .38, *p* = .006, *q* = .02], AF4 [*r*(50) = .38, *p* = .006, *q* = .02], F3 [*r*(50) = .33, *p* = .02, *q* = .04], F4 [*r*(50) = .34, *p* = .02, *q* = .03], F7 [*r*(50) = 0.43, *p* = .002, *q* = .01), F8 [*r*(50) = .34, *p* = .02, *q* = .03] and in the right posterior channels P8 [*r*(50) = .44, *p* = .002, *q* = .01] and O2 [*r*(50) = .34, *p* = .02, *q* = .03]. Significant positive correlations that did not survive the multiple comparison correction were also found in subsets of the same channels across other bands. Specifically, power in the alpha band was positively correlated with the rate of forgetting in channels F8 [*r*(50) = .29, *p* = .04, *q* =.18] and P8 [*r*(50) = .31, *p* = .02, *q* = .18]. Power in the low beta band correlated with the rate of forgetting in channels AF4 [*r*(50) = .30, *p* = .03, *q* = .20], and P8 [*r*(50) = .32, *p* = .02, *q* = .20]. And power in the high beta band correlated with the rate of forgetting in channels AF4 [*r*(50) = .31, *p* = .03, *q* = .14] and F4 [*r*(50) = .29, *p* = .04, *q* = .14]. Finally, and like in all previous analyses, no significant negative correlation was found.

**Figure 7:**
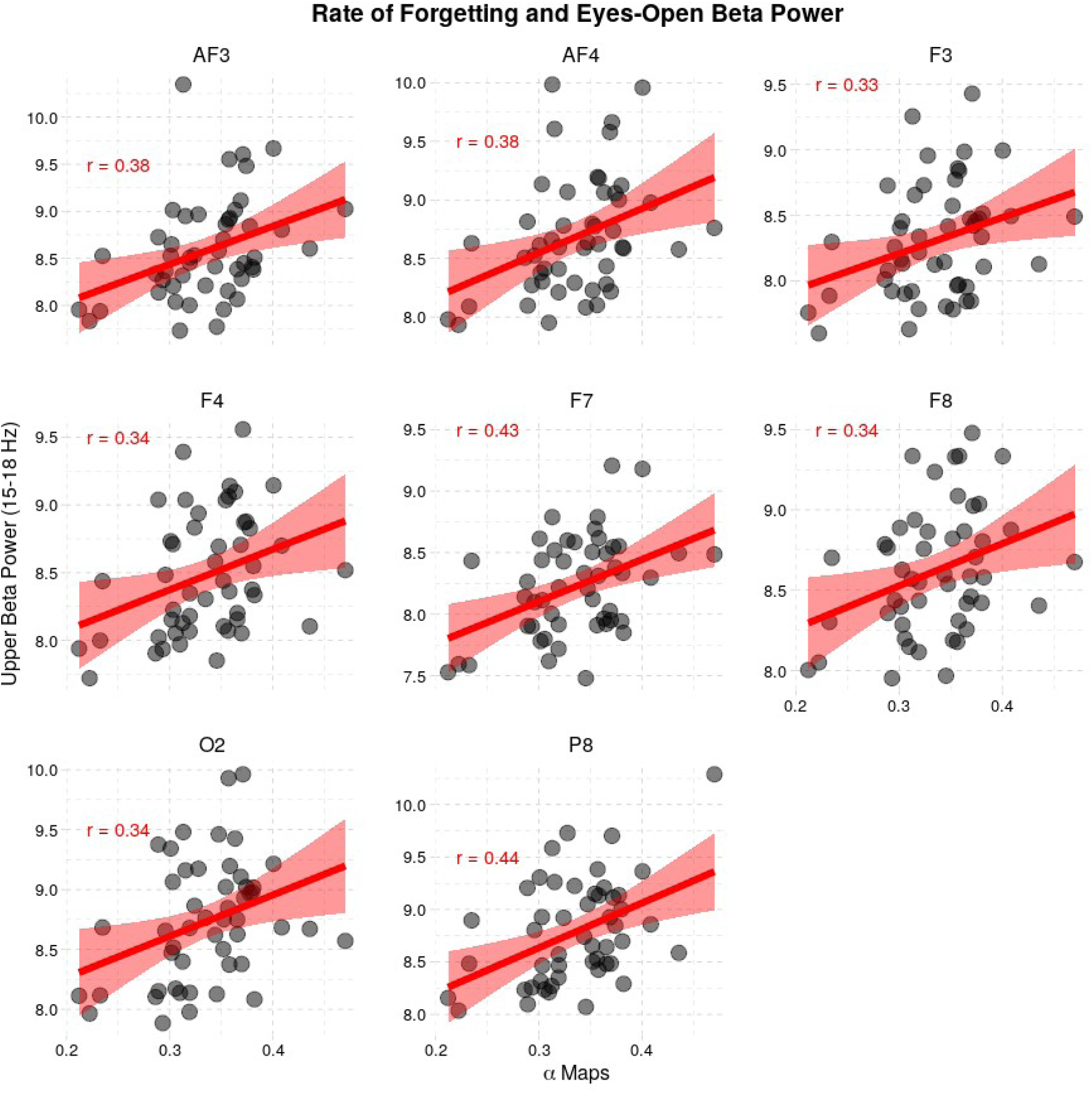
Correlations between power in the upper beta band (15-18 Hz) and the rate of forgetting for visual materials (US Maps) in eyes-open recordings. In all channels, the correlation is significant at q < 0.05 (FDR corrected).

### 3.4 Robust Correlation Analysis

As noted above, the distribution of forgetting rates in our sample is compatible with the distribution reported by previous studies (Sense et al., 2016; 2018; van der Velde et al., 2020). Still, it is worth examining whether the correlations identified in the previous analysis might be primarily driven by a small population of individuals who exhibit very low rates of forgetting. To this end, we re-assessed all of the previous analysis using a robust correlation method developed by Smirnov and Shevlyakov (2014), which greatly reduces the effect of outliers in the data (Shevlyakov & Smirnov, 2011) and is implemented in the package “robcor” for the R statistical language. In general, the robust correlation method reduced the absolute value of most (but not all) correlation coefficients. However, the results did not change significantly in terms of which correlations survived the False Discovery Rate procedure.

The single correlation coefficient that survived FDR correction in the eyes-closed recordings (rate of forgetting for verbal materials, channel P8, low beta band: Fig. 4) was found to be still significant after using a robust correlation algorithm (robust *r* = .36, p = .01), but not to survive FDR correction (*q* = .15). Of the eight correlation coefficients that survived FDR correction in the eyes-open recordings (rate of forgetting for visuospatial materials, channels AF3, AF4, F4, F4, F7, F8, O2, and P8, upper-beta band) all but one (F3) survived error correction when their correlation coefficients were calculated with the robust method (*r* > .32, *p* < .03, *q* < .05). On the other hand, when robust correlations were used, channel AF4 was found to survive FDR correction at high beta as well (*r* = 0.42, *p* = .003, *q* = .03).

Thus, in summary, the application of robust correlation methods did not alter the general pattern of our findings, suggesting that our findings were not driven by few individuals with exceptionally low rates of forgetting.

## 4. Discussion

In this paper, we have provided evidence that model parameters that reliably characterize an individual’s performance can be observed in that individual’s neurophysiological activity at rest. These correlations were found across two sets of behavioral measures (corresponding to the rates of forgetting for verbal and visual materials) and across two different types of EEG recordings (eyes-closed and eyes-open resting-state EEG), and consistently found that spectral power over bilateral frontal and right posterior scalp locations predicted long-term memory forgetting. Importantly, the observed correlations are in the upper range of values that were expected, given the reliability of both behavioral and EEG measures.

It should be noted that brain activity was extracted at rest before participants performed the experimental task. Thus, this finding adds to the growing number of studies showing that task-free resting-state oscillations contain relevant information about an individual’s biology and cognition (Doppelmayr et al., 2002; 2005; Prat et al., 2016). Specifically, this study is the first not only to link a feature of resting-state brain activity to cognition but to a specific parameter in a formal and well-established model.

### 4.1 Differences Across Materials and Recording Modes

Encouragingly, and consistent with the premises of this research, correlations between rates of forgetting and EEG power spectra were found in both verbal and visuospatial materials and in both eyes-closed and eyes-open EEG recordings. That being said, some notable differences remain.

A first notable difference is that verbal and visuospatial materials had different correlations with EEG features. Although the correlation between the rates of forgetting of the two materials were significant (*r* = .59, Figure 1) and the patterns of EEG correlations were similar (*r* = .61, Figure 3; and *r* = .81, Figure 6), they were not identical. The differences between the two could either be due to simple measurement error or to the specific properties of the two materials. Although both hypotheses are possible within the scope of this study, some evidence points to the latter. First, lower the correlation across materials than within the same materials was also found in the original paper by Sense et al (2016). Second, the difference is somewhat reflected in the different topological distributions of the EEG correlations (Figures 3 and 6), with forgetting rate for maps being more often associated with frontal rather than posterior electrodes and with a more bilateral distribution. It remains to be determined, however, whether these differences reflect the representational codes (verbal vs. iconic) of the two materials, or whether they simply reflect a difference in cognitive demands (maps seem harder to process than words, as evidenced by the higher frogetting rate, Figure 1).

It was also the case that the specific recording protocol (eyes-closed vs. eyes-open) yielded differences in which correlations survived correction in the two types of materials to be memorized (verbal vs. visuospatial). Specifically, eyes-open recordings yielded overall better results than eyes-closed recordings, especially in the case of visuospatial materials and especially if robust correlations were used—in which case, no single channel survived the False Discovery Rate correction. As noticed in the Introduction, among the reasons for preferring eyes-open recordings are the fact that they might provide a better baseline for cognitive processes that depend on visual areas, as is likely the case of maps, and for neural processes that occur in the beta band, where most of our effects are found.

Although these differences are worth exploring in future work, we would argue that the totality of the results paints a consistent picture. Figures 3 and 6 show that the patterns of correlations were remarkably similar across eyes-open and eyes-closed EEG, and this similarity suggests that the EEG correlates of rate of forgetting most likely reflect an underlying neural mechanism rather than an intrinsic source of variability that is specific and confounded to that particular way of acquiring data (e.g., individual-specific aspects of visual processing or eye blink artifacts). On the contrary, if the patterns of correlations had been dramatically different between the two types of acquisition, it would have suggested that the correlations were driven by EEG activity that was driven by some spurious characteristic (for example, different noise artifacts due to the different eye movement in the two conditions).

### 4.2 Specificity of Beta Band and Fronto-Parietal Activity to Human Long Term Memory

Although many locations on the scalp were found to show mild correlations with our parameter, the strongest results were consistently found, for both verbal (Swahili words) and spatial material (US maps) and across eyes-open and eyes-closed recordings, in frontoparietal locations (channels AF3, AF4, and P8 in particular) and in a specific frequency band (low to upper beta, 13-18 Hz; Figure 8).

**Figure 8:**
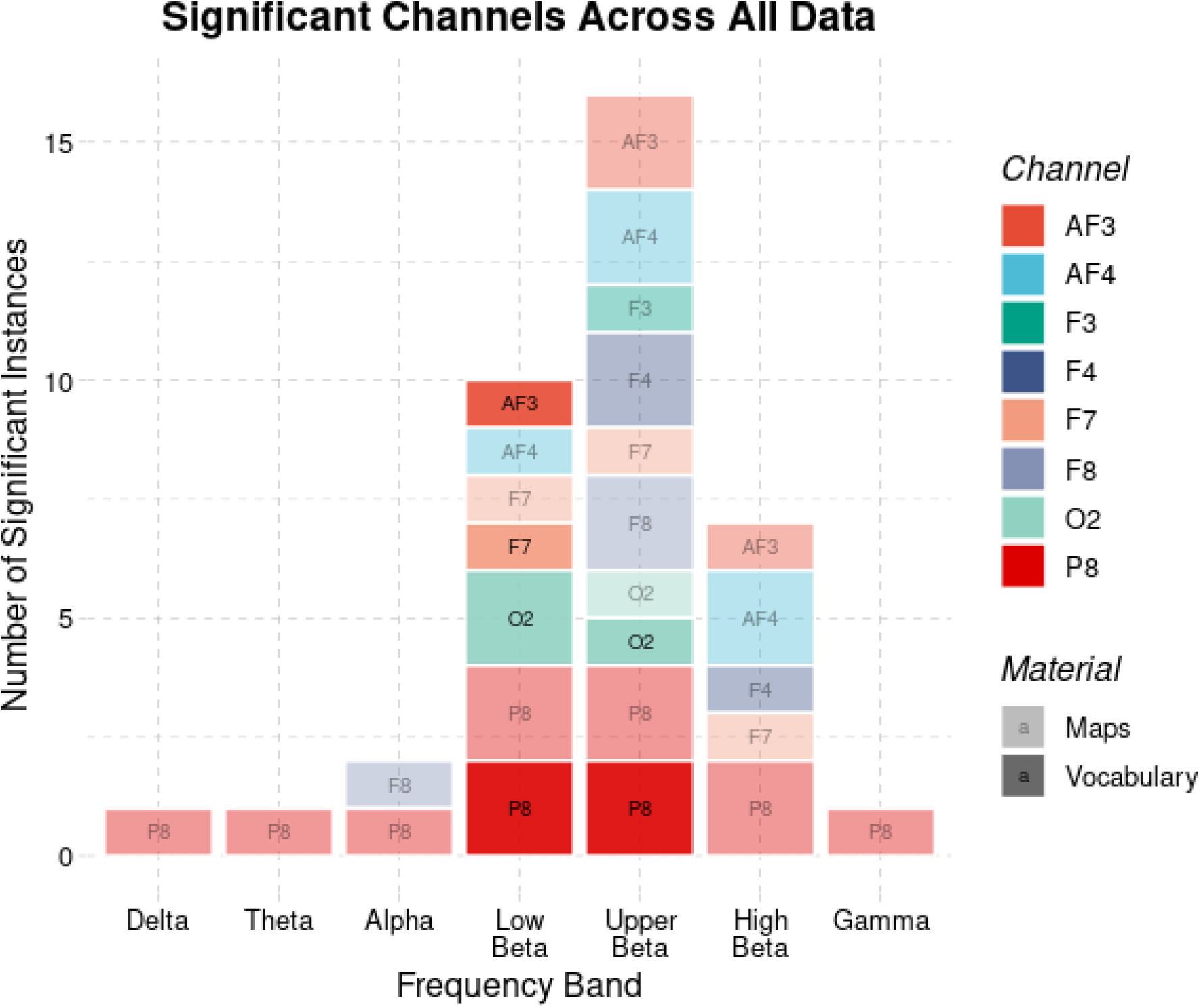
Histogram of significant correlations (p < .05) found across both types of recordings and materials. Full colors represent correlations with Vocabulary; semi-transparent colors represent correlations with Maps. Single-height boxes represent significant results in one recording modality (eyes open or eyes closed), double-height boxes represent significant results in both recordings.

This particular location and frequency band might be unexpected, given that one would expect individual differences in the rate of forgetting to be related to the activity of cortical and subcortical regions in medial *temporal* lobe (Palombo et al., 2018) whose *theta* band activity has been associated with memory formation (Bland & Oddie, 2001; Lega et al., 2012; Solomon et. al., 2019). This discrepancy, however, should not sound surprising. First, the location of an EEG signal over the scalp does not translate directly to an underlying source. Thus, the fact that a signal is recorded over a parietal region does not imply that its source is also parietal. Second, signals from medial regions are notoriously difficult to identify in EEG data, due to their particular position and distance from the scalp (Gavaret et al., 2004; Pizzo et al., 2019). Therefore, it is likely that the signals that are mostly correlated with rates of forgetting do not reflect hippocampal activity per se, but possibly the contribution of other circuits that are ancillary to memory encoding and retrieval and might be indirectly affected by the intrinsic activity of the hippocampus. Specifically, beta power decreases in frontal and parietal regions have been repeatedly found to be associated with deeper encoding and greater retrieval accuracy of the studied material (Hanslmayr et al., 2012; Subramaniam et al., 2019). Furthermore, these fronto-parietal decreases in power, technically known as Event-Related Desynchronizations (ERDs), are also associated with corresponding increases in the fronto-parietal BOLD response (Hanslmayr et al., 2011), which similarly predicts successful encoding and retrieval (Kirchhoff & Buckner, 2006); and with power increases in theta activity in the hippocampus, thus providing a direct link to the neural circuit more directly responsible for memory formation (Sederberg et al., 2007).

The vast majority of the studies relating fronto-parietal beta activity and memory formation, however, have relied on event-related, *task-based* paradigms, while the study described herein employs *task-free*, resting-state recordings. The dissimilarity of the two approaches raises the issue of the relationship between beta power at rest and the amplitude of ERDs in the beta band during a task. A possible connection between the two is that, because higher power at rest indicates greater synchronized and coherent activity of neurons (Pfurtscheller, 1997), it might also limit the ability of these neuronal populations to de-synchronize during memory encoding. Under this hypothesis, populations of fronto-parietal neurons that are spontaneously synchronized (due to a larger number of synapses, for example) have a difficult time to “step out of rhythm” to carry out the computations needed for rich, successful encoding of memories, and their tendency to oscillate together translates into both greater beta power at rest and reduced ERDs during the task—ultimately leading to shallower encoding, greater rate of forgetting, and reduced retrieval accuracy. Thus, our pattern of results is broadly consistent with the known neurophysiology of memory encoding and retention.

### 4.3 Limitations

When interpreting these results, some limitations must be considered. A few of these limitations are intrinsic to the use of EEG: as noted above, it is impossible to precisely identify the source of (resting-state) EEG signals. For this reason, we can only speculate about the brain regions whose activity is associated with the decay rate. By contrast, information about the source would be valuable in understanding what type of information we are dealing with. This limitation could be overcome by using different neuroimaging methods, such as MRI-informed MEG.

The EEG-specific limitations are compounded, in our case, by the use of low-grade systems. While they allowed us to record and collect data quickly and they have been used successfully in research (Jiang et al., 2019; Prat et al., 2016; Zhou et al., 2020), they are also characterized by poor spatial resolution, poor sampling rate, and low sensitivity. Thus, not only is source identification practically impossible, but significant features that are best observed in midline locations (not covered by our setup), or have low amplitude, or occur at higher frequency rates would likely go undetected in this study. However, the use of this system demonstrates that relevant information on individual learners can be obtained using systems that might see usage in class-room settings, opening the door to neuroscience-informed adaptive learning systems.

Also, it is important to notice that, adhering to standards set in similar studies with comparable equipment (Prat et al., 2016, 2019), all of our analyses were restricted to bi-variate correlations. This permits, among other things, an intuitive examination of each correlation in isolation. However, it should be noted that more sophisticated methods, including machine learning approaches that take into account multiple features, could be potentially employed. This would likely result in greater precision at *predicting* specific values of α from EEG data alone, given the fact that the correlations were spread over multiple channels and bands. This would, however, also significantly complicate the interpretation of the results. For the purpose of this study, we prefer to focus on reporting on the existence of multiple electrophysiological signals that correlate with rates of forgetting and their likely neurophysiological origin.

Finally, in this study, brain activity was interpreted in relationship to one specific component of a formal model. Although this model has been successfully employed to estimate the rate of forgetting (Sense et al., 2016; 2018; 2019; Van den Broek et al., 2019), the conceptual framework of which the current model is part of also contains other parameters. For example, the probability of retrieving a memory (for example, the association “kitanda”/”couch”) also depends on spreading activation from the cue (for example, the Swahili word “kitanda”). The amount of spreading activation is assumed to be modulated by attention, and the degree of attention allocated to cues has been found to be predictive of working memory capacity (Lovett, 2001). Thus, the high correlation found in channel P8 may be partially driven by attentional processes that interact with the rate of forgetting during retrieval. The fact that both the location of the channel (right parietal lobe) and the specific band (beta) have been previously connected to the retrieval of verbal material suggests that this might be the case. Future studies in which both factors are parameterized and estimated for each individual would help more precisely identify the nature of the biological processes involved.

### 4.4 Implications for Future Research

These limitations notwithstanding, our results do have significant implications for future research. The first and most obvious is that they outline a method to estimate the value of idiographic trait parameters in the absence of task assumptions and with a very reasonable and cost-effective setup. In our case, for example, the collection of EEG recordings took significantly less time than the collection of behavioral data, even accounting for set-up time. This effect is amplified by the use of consumer-grade headsets, which, while resulting in reduced fidelity recordings, also permit quicker and faster data collection from a larger sample than would be allowed by research-grade equipment. Most importantly, while estimating multiple parameters would require a combination of specific tasks, task-free brain imaging data likely contains signals that reflect multiple parameters of interest. Once collected, every additional parameter can be extracted from the data without increasing the number of sessions or their duration.

In addition to its practicality, the use of task-free neuroimaging data to estimate idiographic parameters provides methodological and conceptual advantages. First, it provides a direct connection to the biological interpretation of a parameter. Second, it provides a useful way to constrain the development of computational cognitive models, as solid correlations with well-established biological correlates could become a litmus test of the construct validity of components/conceptual constructs of computational models.

Of course, for these implications to be realistic, the method we have outlined here would need to be expanded upon and tested using other well-established models and parameters. Nonetheless, we believe in the importance of this approach for future research. As noted in the introduction, idiographic models are necessary for successful translational applications of cognitive research in education and clinical settings, and task-free measures of idiographic parameters provide the most straightforward and biologically-grounded method to tailor models to particular individuals.

## 5. Acknowledgments

All of the supporting data and the analysis code can be found online at https://github.com/UWCCDL/EEG_RateOfForgetting. This research was partially supported by grant FA9550-19-1-0299 from the Air Force Office of Scientific Research to AS.

## Notes

https://github.com/UWCCDL/EEG_RateOfForgetting

https://osf.io/47mu3/

### Competing Interest Statement

The authors have declared no competing interest.

### Summary of Updates

The revised submission has taken into account the feedback of three reviewers. The manuscript has been modified in the following ways: 1. Clarification of the behavioral paradigm. Reviewer 1 pointed out that the description of the adaptive memory task was insufficiently clear. Thus, we have significantly expanded the description of the task in Section 2.3.2 "Swahili Vocabulary Learning Task" and how it relates to the model in Section 2.4.2 "Analysis of Behavioral Data" 2. Additional comparisons across materials and recordings. Reviewers 2 and 3 requested additional comparison across recordings (eyes-open and eyes-closed) and materials (visual s. verbal). Comparisons between the two have been added to all of the Results sections (Sections 3.2.1, 3.2.2, 3.3.1, 3.3.2, and 3.3.3). In addition, a new version of Figure 5 and an entirely new Figure (Figure 8) have been added to facilitate the comparisons. Finally, the revised Discussion now contains a new section (4.1 "Differences Across Materials and Recording Modes") summarizing these differences.

